# Linking Pain and Delirium via Microglial Activation: A Mouse Study Using BSEEG, Behavioral Assays, Immunohistochemistry, and RNA Sequencing

**DOI:** 10.1101/2025.10.29.685463

**Authors:** Kyosuke Yamanishi, Riley Merkel, Nathan James Phuong, Victoria Anderson, Shota Nishitani, Tsuyoshi Nishiguchi, Takaya Ishii, Bun Aoyama, Nipun Gorantla, Hieu Dinh Nguyen, Ilgin Genc, Hisato Matsunaga, Vivianne L. Tawfik, Gen Shinozaki

## Abstract

**Background:** The rising incidence of delirium in surgical and critical care settings, especially in older patients, calls for improved preventative and management strategies. Chronic pain is increasingly recognized as a key risk factor for delirium, and microglial activation may be the central mediator linking these two conditions.

**Methods:** We used a spared nerve injury (SNI) model of persistent neuropathic pain in tandem with a postoperative delirium (POD) mouse model. Pain assessments, electroencephalography (EEG) recording, and immunofluorescence were performed to characterize pain and delirium-like states. Microglia were isolated for RNA-seq to elucidate gene expression changes comprehensively.

**Results:** SNI mice showed persistent mechanical hypersensitivity from Day 7 onwards and demonstrated disrupted sleep-wake patterns in EEG indices. Immunofluorescence revealed sustained microglial activation in both the hippocampus and cortex following SNI. RNA-seq analyses indicated the upregulation of pro-inflammatory pathways (e.g., interleukin-6 production, NF-κB signaling) in SNI mice that also underwent head-mount surgery. Notably, the coincidence of persistent pain and an acute delirium-like state further exacerbated neuroinflammation.

**Conclusion:** Our findings suggest that neuropathic pain-induced microglial activation primes the brain for exaggerated inflammatory responses under additional surgical stress, potentially worsening delirium. Further investigations into microglia-focused therapies may inform novel strategies for mitigating delirium in patients with neuropathic pain.

## 1. Introduction

As population age, an increasing number of patients are developing delirium during the course of critical illnesses or after surgery ^1^. Characterized by acute and fluctuating disturbances in cognition and attention, delirium is also associated with a prolonged hospital stays and poorer clinical outcomes^2-5^. Consequently, there is an urgent need for effective strategies to prevent and manage delirium. Among the known risk factors, inadequate pain control, particularly in patients with severe pain, has been identified as a key contributors to delirium^6^. In clinical practice, the importance of appropriate pain management for delirium prevention is widely recognized^6^.

The activation of microglia, immune cells in the central nervous system, appears to play a major role in the pathophysiology of both pain and delirium ^1, 7, 8^. Recent evidence suggests that pain signals, originating from peripheral nerves, may activate microglia worsen anxiety and depression^9, 10^. Inflammatory processes in the brain involving microglial activation by peripheral insults have also been implicated in the pathogenesis of delirium. It is, therefore, hypothesized that microglia may serve as a key mediator linking pain and delirium^11, 12^.

Based on these previous reports, the present study tests the hypothesis that microglia mediate the interaction between pain and delirium. Specifically, we used the spared nerve injury (SNI) mouse model, a well-established model of post-surgical pain/nerve injury-induced pain, in tandem with our previously developed postoperative delirium (POD) mouse model which is quantified using the bispectral EEG (BSEEG) method^13, 14^. In this POD model, EEG head-mount implantation surgery itself is used as the surgical insult, inducing postoperative EEG changes along with deficits in attention and cognitive function^14^. By performing an SNI procedure to induce a persistent painful state and subsequently modeling POD, we aim to investigate the mechanisms of delirium onset under pain conditions and to evaluate how microglial activation contributes to such interaction.

## 2. Materials and Methods

### Animals

Male C57BL/6J mice (7–8 weeks old) were housed under a standard 12-hour light/dark cycle with free access to food and water at a temperature of 23–25 °C and humidity of 40%–50%. The animal experiments were conducted following a protocol approved by Stanford’s Administrative Panel on Laboratory Animal Care (APLAC), which is accredited by the Association for Assessment and Accreditation of Laboratory Animal Care. (approval number: APLAC-34459) In this study, mice were divided into four experimental groups. As shown in Figure 1A, SNI group mice underwent spared nerve injury (SNI) surgery on Day 0, while control mice were only exposed to anesthesia exposure. One week after SNI surgery, head-mount surgery was performed to induce a delirium model, whereas the control group underwent anesthesia exposure only. Accordingly, the four groups were: (1) SNI surgery + head-mount surgery group (SNI+HM group), (2) SNI surgery + No head mount surgery group (SNI+NoHM group), (3) No SNI surgery + head-mount surgery group (NoSNI+HM group), and (4) No SNI surgery + No head-mount surgery group (NoSNI+NoHM group).

**Figure 1.**
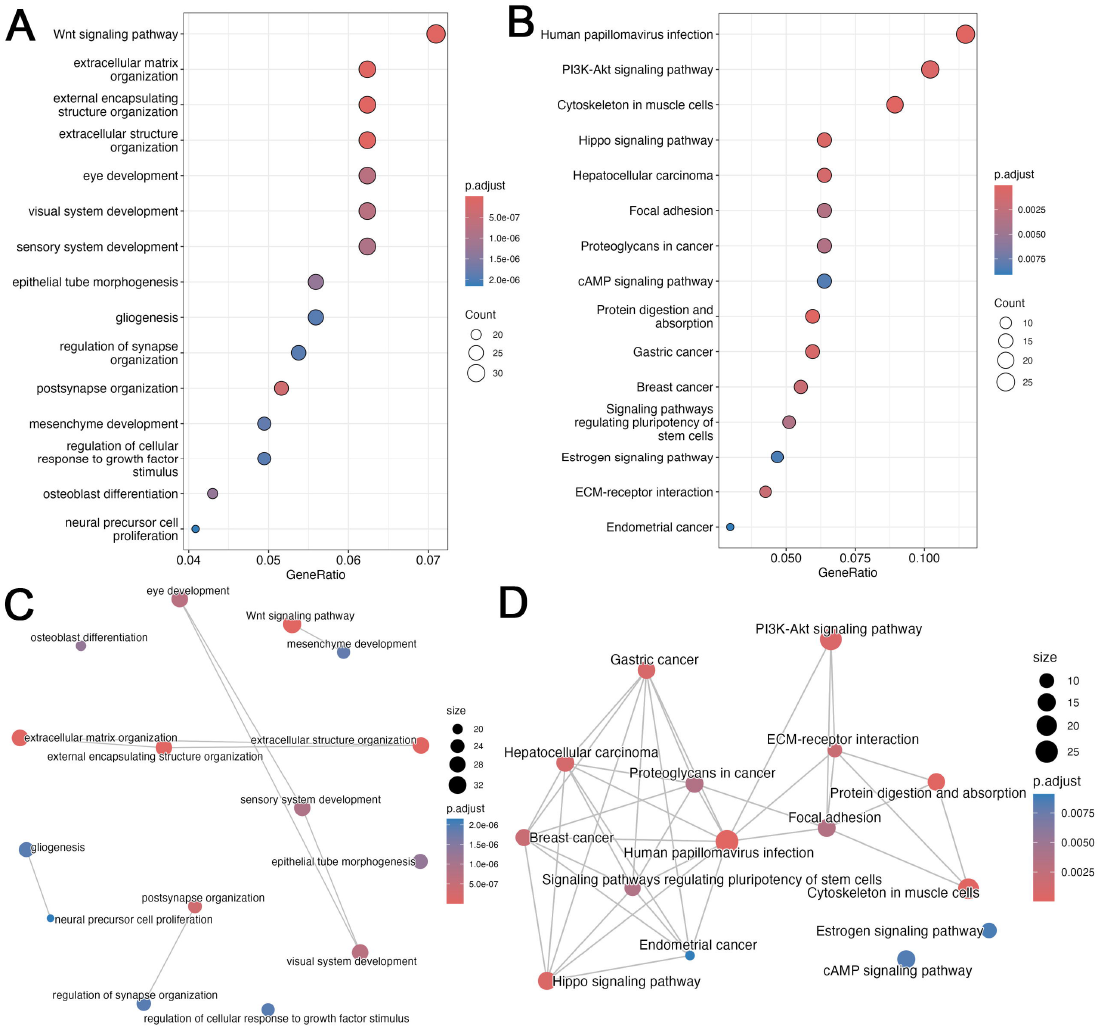
SNI-induced mechanical hypersensitivity and changes in EEG parameters. (A) Experimental timeline. On Day 0, mice underwent either SNI surgery or not. The von Frey testing was performed on the indicated days following surgery. (B) Time-course changes in mechanical threshold measured by the von Frey filament test. The SNI group (orange) shows significantly decreased thresholds compared to the control group (blue) after surgery. (C) Changes in sBSEEG over time for the control (blue) and SNI (orange) groups. (D) Wave frequency in the 10 Hz range, shown by daily measures (left) and mean values in first four days (right). (E) Power in the 3 Hz (delta) range, shown as daily measures (left) and mean values in first four days (right). All error bars represent the standard error of the mean (SEM). *p < 0.05, ***p < 0.001. (B; *n*=5 per group, C-E: *n* = 8 each group)

### EEG Electrode Head-Mount Placement Surgery

EEG electrode head mount placement surgery was performed as in our previous studies^15, 16^. Briefly, the surgery was performed under isoflurane anesthesia (1–3% inhalation). The skull was exposed, and the head mount (#8201, Pinnacle Technology, Inc., Lawrence, KS) was positioned so that its center aligned with the midline of the skull; the mount’s anterior holes were located 2 mm anterior to bregma, and the posterior holes 2 mm anterior to lambda (±2 mm mediolateral). Four each of the four holes in the head mount, a 23-gauge needle was used to bore corresponding holes in the skull. Stainless-steel screws (#8209 for anterior, #8212 for posterior; Pinnacle Technology, Inc.) were manually inserted into each hole. The head mount was then secured with dental cement. Postoperative analgesia was provided via subcutaneous meloxicam injection (5 mg/kg).

### EEG Recording and EEG Signal Processing for BSEEG Scores

The mice were tethered to an EEG system (#8200-K1-SL, Pinnacle Technology, Inc.) for continuous data collection throughout the remainder of the experiment. EEG signals were recorded from the right frontal cortex using the right anterior skull screw (EEG2) and from the right parietal cortex using the right posterior skull screw (EEG1). The left frontal screw was designated as ground and the left parietal screw served as reference. Based on our previous findings indicating higher sensitivity of EEG2 over EEG1, the EEG2 channel was primarily used for BSEEG scoring^14^.

Raw EEG data were converted into power spectral densities to obtain BSEEG scores as previously described^14^. Signal acquisition was performed using Sirenia software, recordings were exported as EDF files. The resulting files were subsequently processed through a web-based analysis tool (https://sleephr.jp/gen/mice7days/) that automatically calculates the ratio of 3–10 Hz power, defined as the BSEEG score.

To highlight the differences in BSEEG scores relative to the postoperative steady state, we defined the standardized BSEEG (sBSEEG) score as the difference between daytime/night BSEEG scores on each postoperative day and the daytime BSEEG scores on postoperative Day 7, which was set as sBSEEG score = 0. This approach was adopted because baseline EEG cannot be recorded prior to head-mount surgery, and young mice are generally considered fully recovered by Day 7 based on our previous data^14^. Thus, most mice were estimated to have fully recovered and reached a steady state in their BSEEG scores by postoperative Day 7.

### EEG Analysis of 10Hz and Power Spectrum Focusing on the Delta

In this study, we used fast Fourier transform (FFT) to calculate the power spectral density (PSD) in order to evaluate the frequency characteristics of electroencephalography (EEG). Specifically, we divided the EEG data into 10-minute segments, performed FFT on each segment, and then averaged the resulting PSD values to obtain the power in specific frequency bands. We focused on two frequency ranges: the 3 Hz and the 10 Hz band.

First, for the 3 Hz band, PSD values within 3 Hz were averaged for each 10-minute segment to define the delta power. The 10 Hz band has been reported to exhibit high specificity for the brain during quiet waking^17-19^. So, we used the PSD in the 10 Hz range as an index of waking activity. Finally, we investigated how the calculated 3 Hz and 10 Hz frequencies relate to sleep/wake stages and potential changes in sleep structure.

### Spared Nerve Injury

SNI surgery was performed as previously described^13^. Briefly, mice were anesthetized using isoflurane (Induction: 3-3.5%, Maintenance: 1.5%), and a small incision was made on the left thigh. Blunt dissection was performed through the biceps femoris muscle to expose the sciatic nerve and its three branches: the common peroneal, tibial, and sural nerves. The common peroneal and tibial nerves were ligated with a 5–0 nylon suture (ETHILON™ ref#1668G) and subsequently axotomized using small spring scissors. The sural nerve was preserved as the “spared nerve.” The incision was then closed using surgical staples, which were removed 7 days after the SNI surgery.

### Von Frey Test

To assess mechanical sensitivity in mice, we employed the widely validated von Frey “up-down” method. This method utilizes a set of 10 von Frey filaments (Stoelting Co. Wood Dale, IL, USA), that exert logarithmically increasing forces ranging from 0.02 to 4.0 grams (specifically, 0.02, 0.04, 0.07, 0.16, 0.4, 0.6, 1, 2, and 4 g). The filaments are applied perpendicularly to the plantar surface of the hind paw with sufficient force to induce a slight bending of the filament. A positive response is defined as a rapid withdrawal of the paw from the filament stimulus within 4 seconds. Using the up-down statistical method as described previously, we calculated the 50% withdrawal mechanical threshold scores for each mouse^20^. Following the initial positive response, four additional responses were recorded, resulting in a total of five withdrawal responses per mouse during each testing session.

### Immunofluorescence Staining

Immunofluorescence staining was conducted to assess differences in neuroinflammation. Mice were deeply anesthetized with 4% isoflurane and underwent transcardial perfusion with 20 mL of saline followed by 4% periodate-lysine-paraformaldehyde fixative. After fixation, the brains were extracted, and coronal brain sections 40 µm thick, which included the dentate gyrus of the hippocampus, were prepared. These sections were blocked with 5% bovine serum albumin for 1 hour. Subsequently, the sections were incubated overnight at 4°C with the following antibodies: anti-CD68 (1:400, cat. No. #97778S, Cell Signaling Technology, Inc., Danvers, MA, USA) and anti-IBA1 (1:400, cat. No. #019-19741, FUJIFILM Wako Pure Chemical Corporation, Osaka, Japan). Next, fluorochrome-conjugated secondary antibodies were applied for 2 hours at room temperature: donkey anti-Goat IgG (H+L) Alexa Fluor® 488-conjugated (1:500 dilution; cat. no. A11055; Thermo Fisher Scientific, Waltham, MA, USA). or goat anti-rabbit IgG H&L Alexa Fluor® 594-conjugated (dilution 1:500, cat. No. A32740; Thermo Fisher Scientific Inc.). The sections were then mounted with Vectashield containing 4,6-diamidino-2-phenylindole (DAPI) (cat. No. H-1800; Vector Laboratories, Inc., Burlingame, CA, USA) and visualized by a microscope (BZ-X800, KEYENCE Corp., Itasca, IL, USA). Images were scanned and analyzed using BZ-X800 viewer and BZ-X800 Analyzer software (KEYENCE Corp.), and the number of positive cells along with their density was quantified and averaged across 8-10 sections per mouse, as previously described^21, 22^.

### Microglia Isolation

Initially, the brain was perfused with PBS containing 0.1 mM EDTA to facilitate extraction. The tissue was then dissociated using the Adult Brain Dissociation Kit (130-107-677, Miltenyi Biotec, Inc., MD, USA) to isolate brain cells following the manufacturer’s protocol. Subsequently, CD11b-PE antibody (130-113-806, Miltenyi Biotec) was applied to label the cells, and Ultrapure-PE-microbeads (130-105-639, Miltenyi Biotec) were used to specifically bind the antibody-labeled cells. Magnetic Activated Cell Sorting (MACS) was then employed to isolate the CD11b-positive cells, allowing for the sampling of microglia.

### Sample Preparation and Quality Control

Total RNA was isolated from each sample with the RNeasy Micro Kit using manufacturer’s protocols (74004, QIAGEN, Hilden, Germany). RNA integrity and concentration were assessed with an Agilent 4150 Bioanalyzer (Agilent Technologies). Only samples with RNA integrity number (RIN) values above the acceptable threshold (RIN ≥ 8) were retained for downstream library construction.

### RNA Sequencing

mRNA libraries were prepared from total RNA by poly(A)-enrichment (or ribodepletion, if applicable). The fragmented mRNA (or RNA) was reverse-transcribed into cDNA, followed by second-strand synthesis. After end repair, adapter ligation, and size selection, libraries were PCR-amplified. Each library was validated for size and concentration (Qubit). Equimolar pools were then loaded onto an Illumina sequencer, producing paired-end reads stored in FASTQ format. Raw reads in FASTQ format were examined with tools such as fastp to remove adapters and low-quality bases. Reads passing quality thresholds (Q20, Q30, and GC content) were carried forward. Clean reads were aligned to the reference genome (mm39) using HISAT2, which utilizes global and local FM indexes for spliced read alignment. Alignment statistics were summarized. Mapped reads were assigned to genomic features using featureCounts^23^. Normalized expression levels were calculated (FPKM) and assessed via box plots or density plots to ensure sample consistency. Details of RNA-seq can be found on the Gene Expression Omnibus website (GSE308437).

### Differential Gene Expression Analysis

Raw counts from featureCounts were analyzed in DESeq2 and edgeR, employing negative binomial models for normalization and dispersion estimation^24^. Genes with P ≤ 0.05 were considered significant. Volcano plots, heatmaps, and MA plots were used to visualize these results as previously shown^21^.

### Functional Enrichment Analysis

Differentially expressed genes were assessed for Gene Ontology (GO) term enrichment and Kyoto Encyclopedia of Genes and Genomes (KEGG) pathway overrepresentation^25^. In this study, the 500 genes with the smallest P values from the differential expression analysis were selected for subsequent functional enrichment analyses^26^. We used edgeR for enrichment tests, and the top enriched GO terms or KEGG pathways were visualized in bar plots or bubble plots.

### Statistical Analysis

All results are expressed as mean ± standard error of the mean and were analyzed using Sigmaplot™ software (version 11.0; Systat Software Inc., SanJose, CA, USA). Immunofluorescence, von Frey test, and EEG data were analyzed by two way-ANOVA first, and if significance was found, the Holm-Sidak or Tukey post-hoc test was applied. For statistical analysis of RNA-seq, a Student’s t-test or Mann–Whitney U test was applied. Differences were considered statistically significant when p < 0.05.

## 3. Results

### Postoperative time-course changes in an SNI model: behavioral and physiological assessment parameters

The experimental timeline is shown in Figure 1A, with surgery performed on Day□0 and subsequent behavioral measurements obtained on Days□7, 9, 14, and 21. The mechanical withdrawal threshold was markedly decreased in the SNI group compared to the no-SNI (control) group from Day□7 onward, indicating successful induction of neuropathic pain (Figure 1B). In Figure□1C, the sBSEEG score was significantly lower in the SNI group. Because our previous studies have shown that BSEEG increases in young mice during the first four days post-surgery^14^, we calculated the average power of the 10 Hz and 3 Hz frequency band during the daytime and nighttime over the first four days post-surgery. The results are shown on the right sides of Figures 1D and 1E. While there was no significant difference in 3 Hz frequency, a significant increase in the 10 Hz frequency was observed in the SNI group (Figure□1D, E).

### SNI induces sustained microglial activation in both hippocampus and cortex

We compared microglial activation in the hippocampus (Figure 2A-C) and cerebral cortex (Figure 2D-F) between the control and SNI groups. Representative immunofluorescence images (Figure 2A, 2D) showed a marked increase in number of cells positive for IBA1 (green) and CD68 (red) in the SNI group compared to the control. The quantitative measurements reveal that the numbers of both IBA1-positive cells (Figure 2B, 2E) and CD68-positive cells (Figure 2C, 2F) are significantly higher in the SNI group at all examined time points. In the hippocampus (Figure 2B-C), the elevation in microglial markers becomes apparent by Day 1 post-surgery and remains elevated through later time points, indicating ongoing microglial activation. Similarly, in the cortex (Figure 2E-F), a significant increase in microglial marker expression is observed from the early postoperative phase and persists over time.

**Figure 2.**
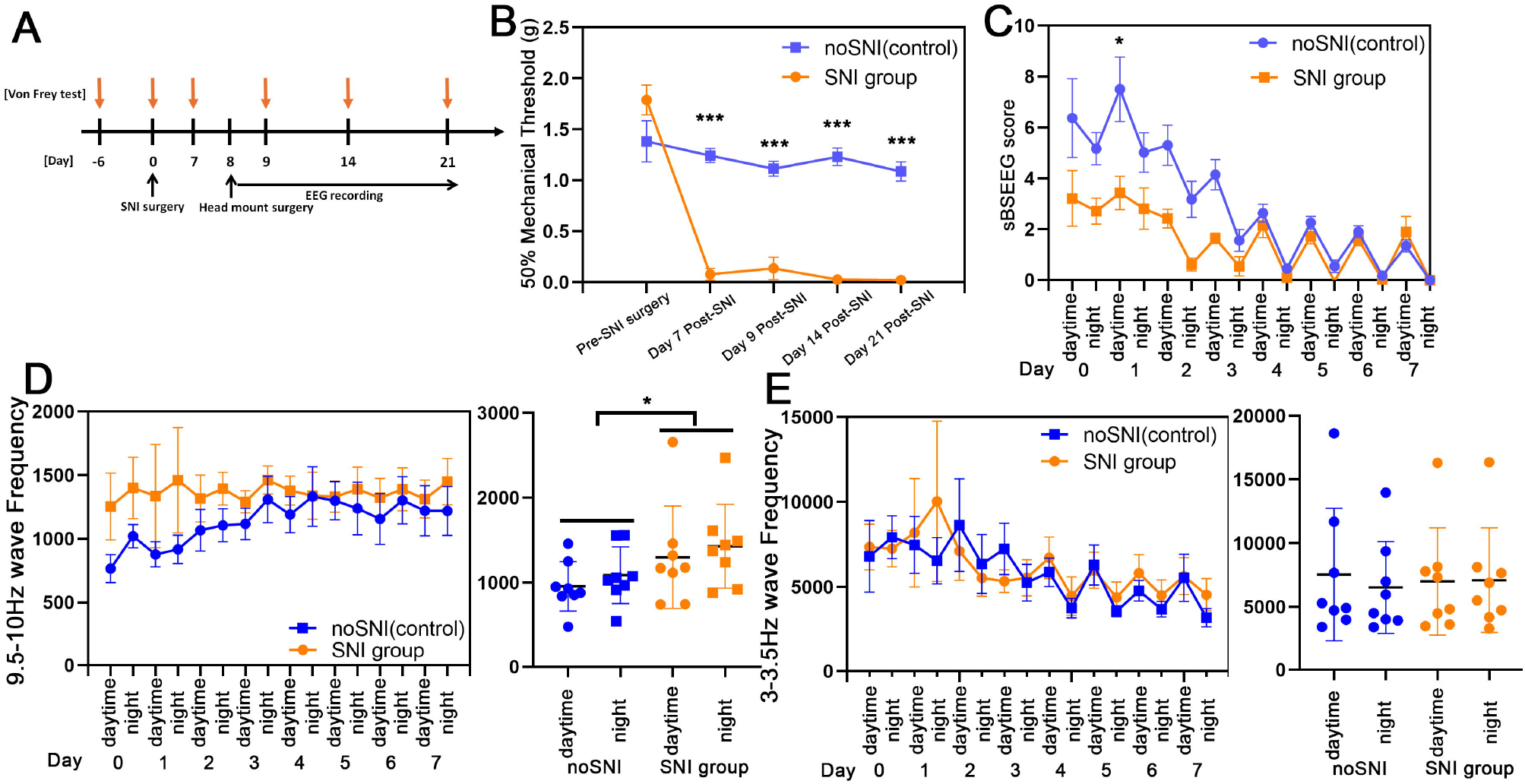
Comparison of microglial activation in the hippocampus and cortex. (A, D) Representative immunofluorescence images of IBA1 (green) and CD68 (red), with DAPI (blue) for nuclear staining. The upper panels, noSNI, show tissue from mice without SNI surgery, and the lower panels, SNI, show tissue from mice sacrificed at the indicated days after head mount (HM) surgery and SNI surgery. Changes in IBA1- and CD68-positive cells were observed in both the hippocampus (A) and cortex (D), respectively. Scale bar = 200 µm. (B, C, E, F) Quantification of the number of IBA1-positive (B, E) and CD68-positive signals (C, F) in the hippocampus (B, C) and cortex (E, F). Values are expressed as means ± SEM. *p < 0.05, **p < 0.01, ***p < 0.001, *n* = 4 per each group.

### 3.3. RNA-Seq

#### Principal component analysis (PCA)

Initially, Principal component analysis (PCA) of the gene expression profiles clearly distinguished the sample groups, indicating substantial transcriptional divergence among them (Figure 3A).

**Figure 3.**
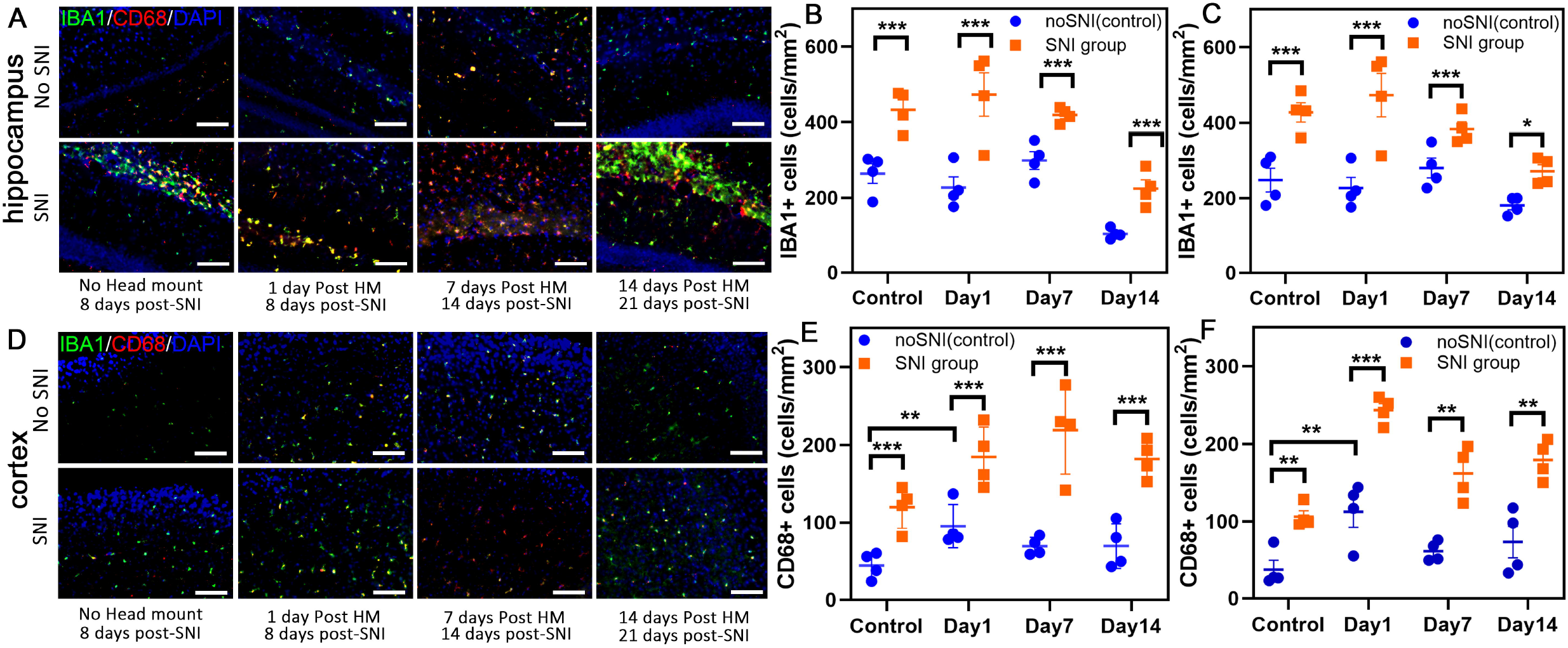
Transcriptomic analysis and functional enrichment of differentially expressed genes (DEGs). (SNI+HM group vs NoSNI+HM group) (A) Principal component analysis (PCA) plot illustrating the clustering of samples based on gene expression profiles. Each dot represents an individual sample, colored according to experimental group. (B, C) Dot plots showing Gene Ontology (GO) biological processes (B) and Kyoto Encyclopedia of Genes and Genomes (KEGG) pathway enrichment (C) analyses of DEGs. Dot size indicates the number of genes enriched in each term, and color gradient represents the adjusted p-value (q-value). (D, E) Network plots visualizing the relationships among enriched GO biological processes (D) and KEGG pathways (E). Node size corresponds to the number of genes involved, and node color indicates the adjusted p-value.

#### Influence of SNI among HM groups (SNI+HM vs NoSNI+HM)

First, we investigated how preexisting pain influences microglial gene expression following head-mount surgery by comparing the SNI+HM group to the NoSNI+HM group. We performed Gene Ontology (GO) enrichment analysis on the identified set of genes (Supplementary Table 1). As shown in Figure 3B, the results showed significant enrichment in pathways related to inflammatory responses including “interleukin-6 production (GO:0032635),” “negative regulation of cytokine production (GO:0001818),” “tumor necrosis factor superfamily cytokine production (GO:0071706),” and “response to lipopolysaccharide (GO:0032496, GO:0071222)”. Subsequently, KEGG pathway analysis revealed significant enrichment in “Apoptosis (mmu04210),” and the “NF-κB signaling pathway (mmu04064)”. These enrichments were also associated with regulating inflammation (Figure 3C).

Network analysis demonstrated that the enriched biological processes are interconnected and form distinct clusters, particularly in areas concerning reactive oxygen species (ROS) responses and inflammation/immune regulation (Figure 3D). KEGG pathway analysis consistently identified not only inflammation-related signaling but also other signaling such as hormone and calcium/cAMP/cGMP signaling as significantly enriched (Figure 3E) (Supplementary Table 2).

#### Influence of SNI among NoHM groups (SNI+NoHM vs NoSNI+NoHM)

Next, in order to investigate the impact of SNI alone on microglia, we extracted microglia from the SNI+NoHM and NoSNI+NoHM group, and performed gene expression analysis.

GO analysis revealed significant enrichment in several biological processes related to extracellular structures and development (Figure 4A). The pathways associated with “Wnt signaling,” “postsynapse organization,” and “neural precursor cell proliferation,” were found in GO analysis. However, few immune-related GO terms were detected, suggesting minimal involvement of immune-specific processes in this context (Supplementary Table 3).

**Figure 4.**
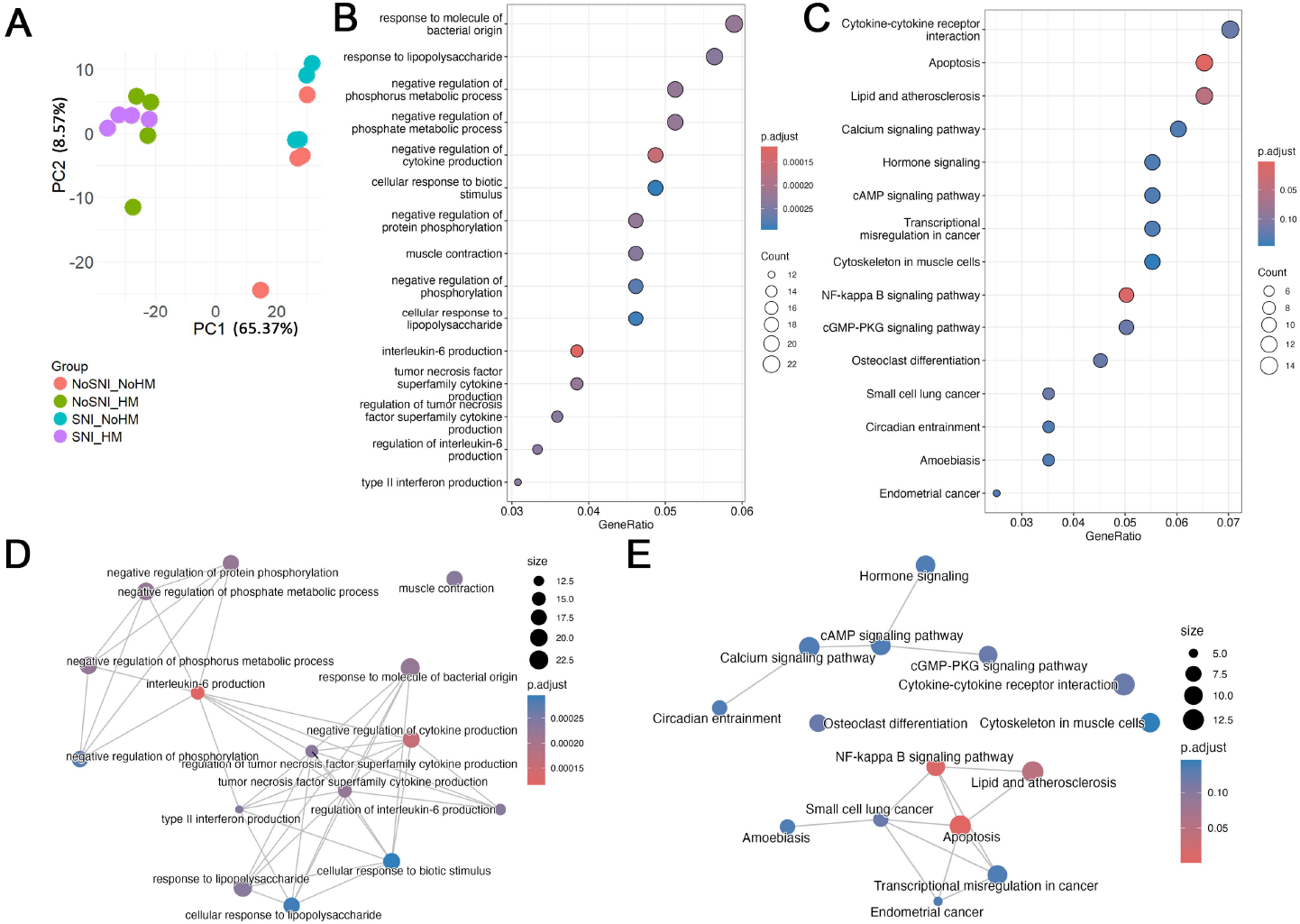
Functional enrichment analysis and pathway interaction networks. (SNI+NoHM group vs NoSNI+NoHM group) (A, B) Dot plots illustrating significantly enriched biological processes (A) and signaling pathways (B). Dot size represents the number of genes involved (gene count), and color gradient indicates the adjusted p-value (q-value), with red indicating higher significance. (C, D) Network diagrams depicting interactions among enriched biological processes (C) and signaling pathways (D).

KEGG pathway analysis identified considerable enrichment in diverse pathways, including those related to core regulatory mechanisms (e.g., PI3K-Akt signaling pathway and cAMP signaling pathway) (Figure 4B). Additionally, “ECM-receptor interactions” which govern cell adhesion and extracellular matrix remodeling, were enriched. As with the GO analysis, immune-related pathways were not detected (Supplementary Table 4).

Network analysis of enriched GO terms revealed that extracellular matrix organization and epithelial morphogenesis play central roles (Figure 4C). KEGG pathway network analysis indicated that the PI3K-Akt signaling pathway occupies a central position, closely interacting with non-immunological functions (Figure 4D).

## 4. Discussion

Considering the growing incidence of delirium among older adults undergoing surgery or intensive care, as well as the recognition that inadequate pain management is one of its risk factors, we conducted this study using a spared nerve injury (SNI) model, which is known to induce a sustained pain-like state, together with a delirium model, quantified by the BSEEG. We aimed to investigate how microglial activation might link pain and delirium. Our results demonstrated that 1) postoperative pain induced by SNI exacerbated microglial activation confirmed by immunofluorescence staining and RNA-seq; 2) EEG analyses revealed transient disruptions in sleep-related indices in SNI mice, while behavioral assessments confirmed that SNI mice experienced mechanical pain hypersensitivity.

In our SNI model, not only was a prolonged reduction in mechanical threshold (i.e., mechanical hypersensitivity in Figure 1B) observed following peripheral nerve injury, but EEG analysis also suggested disruptions in sleep-related indices (Figure 1C-1E). However, while pain behaviors were persistent, the changes in BSEEG were transient.

Clinically, it is well recognized that severe pain or inadequate pain control can be a risk factor for delirium^6, 27^. In this study, we used our POD mouse model, in which we’ve previously observed a postoperative increase in sBSEEG score^14, 28, 29^. Contrary to our hypothesis, however, sBSEEG scores were lower in the SNI model than non-SNI control (Figure 1C). To further investigate this result, we performed an additional EEG analysis, which revealed an upward trend in 10Hz frequency power and no postoperative change in 3 Hz frequency power (Figures 1D and 1E). Notably, the 10Hz frequency band used to calculate the BSEEG in mice showed an increase in the SNI model (as shown in Figure 1D), which is thought to reflect pain-induced hyperarousal, consistent with previous research^30^. Notably, whether or not SNI surgery was performed, the circadian (day-night) rhythm returned to normal after postoperative day 4, underscoring that our model adequately recapitulates the transient nature of delirium^15^. These findings suggest that, in mice subjected to SNI surgery, a hyperarousal state induced by pain coexists with a delirium-like condition coexists.

Previous research reported increased numbers of microglia in the spinal cord following SNI surgery^31^. In the present study, we observed a marked and sustained elevation of microglial markers (IBA1, CD68) in the hippocampus and cortex after SNI surgery. Even one-week post-SNI without the head-mount surgery, the numbers of activated microglia increased significantly. This result suggests that when pain signals originating from peripheral nerves reach the central nervous system, they activate microglia in higher-order brain regions such as the hippocampus and cerebral cortex. Of note, head-mount surgery one week following SNI resulted in a further significant rise in microglia including activated forms in the hippocampus and cortex (Figure 2). Prior studies have similarly indicated that inflammatory responses within the central nervous system following peripheral insults such as surgery play a critical role in delirium^32, 33^. Thus, our present data suggests a potential mechanism linking delirium and pain wherein an already heightened microglial activation induced by pain can further exacerbate neuroinflammation, ultimately intensifying disruptions in brain functions such as delirium.

According to previous studies, delirium is associated with neuroinflammation involving microglia, tumor necrosis factor (TNF)-alpha, and IL-6^32, 34, 35^. In a mouse model of delirium, IL6 mediates the delirium-like phenotypes and IL6 can be produced by activated microglia^36, 37^. Our RNA-seq analysis of microglial gene expression following head-mount surgery showed remarkable changes in inflammatory pathways (e.g., “interleukin-6 production [GO:0032635]” and “tumor necrosis factor superfamily cytokine production [GO:0071706]”) that differed significantly depending on whether SNI surgery had been performed. These results align well with our findings demonstrating that when a delirium-like state is induced in addition to the pain, neuroinflammation is further exacerbated.

This study includes a few limitations. 1) Employing older or female mice, as well as comorbidity models, would better reflect clinically vulnerable populations and enhance generalizability of our findings. 2) Additional EEG analyses and more precise behavioral measures are needed to capture the acute and reversible nature of delirium. In particular, reliable indices for altered consciousness and attention deficits in mouse models would be informative. 3)While regulating microglial activity may mitigate both pain and delirium, broader approaches such as preserving synaptic function should be also explored.

In conclusion, our findings demonstrate that chronic pain, modeled by SNI, not only exacerbates postoperative delirium-like change but also intensifies neuroinflammatory responses, particularly through microglial activation. By integrating behavioral pain assessments, EEG analyses, histological observations using fluorescent immunostaining, and transcriptomic profiling of microglia, we revealed how persistent neuropathic pain primes the central nervous system for inflammation and potentially heightens susceptibility to delirium. These data highlight the importance of early detection and control of pain followed to reduce the risk of delirium in vulnerable surgical populations. Future studies involving older and comorbid mouse models, as well as more nuanced measures for delirium-like states, will be important in translating these discoveries into clinically effective strategies.

## Declarations

### Ethical Approval and Consent to participate

The animal experiments were conducted following a protocol approved by an Institutional Animal Care and Use Committee. At Stanford, the Institutional Animal Care and Use Committee is known as Stanford’s Administrative Panel on Laboratory Animal Care. Stanford’s Administrative Panel on Laboratory Animal Care is accredited by the Association for Assessment and Accreditation of Laboratory Animal Care. (Approved number: APLAC-34459)

### Consent for publication

Not applicable.

### Availability of data and materials

Details of RNA-seq can be found on the Gene Expression Omnibus website (GSE308437). The datasets shown and/or analyzed in the present study are available from the corresponding author upon reasonable request.

### Authors’ contributions

Gen Shinozaki conceptualized and coordinated the entire research study. Kyosuke Yamanishi designed and conducted experiments, acquired data, analyzed data, and wrote the initial draft of the manuscript. Riley Merkel, Nathan Jame Phung, Victoria Anderson, and Shota Nishitani conducted experiments and analyzed data. Tsuyoshi Nishiguchi, Nipun Gorantla, Takaya Ishii, Bun Aoyama, Nipun Gorantla, Hieu Dinh Nguyen, Ilgin Genc, Vivianne L. Tawfik, and Hisato Matsunaga critically reviewed the manuscript. Kyosuke Yamanishi and Gen Shinozaki wrote the final version of the manuscript.

## Acknowledgements

We thank the Stanford Animal Care Facility staff for their technical support.

## Supplementary Table Captions

**Supplementary Table 1.**

Gene Ontology (GO) enrichment results for microglial DEGs (SNI+HM vs NoSNI+HM).

**Supplementary Table 2.**

KEGG pathway enrichment results for microglial DEGs (SNI+HM vs NoSNI+HM).

**Supplementary Table 3.**

Gene Ontology (GO) enrichment results for microglial DEGs (SNI+NoHM vs NoSNI+NoHM).

**Supplementary Table 4.**

KEGG pathway enrichment results for microglial DEGs (SNI+NoHM vs NoSNI+NoHM).

